# Stress-induced motivational impairment is marked by diminished frontocortical cellular communication and neuropeptide signaling

**DOI:** 10.64898/2026.07.11.737837

**Authors:** Puja Parekh, Devin Rocks, Margaux Kenwood, Jacob Roshgadol, Hermany Munguba, Conor Liston

**Affiliations:** Department of Psychiatry, Weill Cornell Medicine, New York City, NY 10021; Department of Neuroscience, The University of Texas at Dallas, Richardson, TX 75080; Department of Biomedical Engineering, University of California, Davis, CA; Department of Neuroscience, Physiology and Pharmacology, University College London, London, UK

**Keywords:** Stress, anterior cingulate cortex, motivation, spatial transcriptomics, cell-cell communication

## Abstract

**Background:** Repeated stress is a risk factor for developing motivational deficits which are common across a variety of disease states including depression and are particularly resistant to treatment with conventional pharmacotherapies. Amotivation is multifaceted and can be caused by impairments in value learning, reward anticipation, and cost-benefit decision-making. Importantly, not all individuals who experience chronic stress develop motivational symptoms, suggesting there may be neurobiological signatures of resilience.

**Methods:** We developed a novel head-restrained effortful reinforcement task in which anticipatory and consummatory behavior can be tracked. Chronic non-discriminatory social defeat stress combined with behavioral analysis and spatially resolved RNA sequencing were used to determine the transcriptional signatures of stress in the anterior cingulate cortex of mice with varying levels of motivational impairment as well as unstressed controls.

**Results:** While stress led to a general impairment in effortful reward seeking, animals differed in the extent of behavioral deficit, with increased ‘susceptibility’ marked by a unique set of differentially expressed genes within the anterior cingulate cortex (ACC). By leveraging the spatial component of our data, we were further able to identify altered interactions from inhibitory neurons and astrocytes to excitatory pyramidal cells, which correlated with intact or impaired motivated responding following stress exposure.

**Conclusions:** Chronic psychosocial stress results in divergent effects on motivated behavior and distinct ACC transcriptional signatures that are concentrated in excitatory pyramidal neurons. Cell interaction analysis implicates enhanced inhibitory neuropeptide signaling and reduced astrocytic contact signaling as upstream markers of motivational resilience and point toward ACC hyperexcitability as a targetable feature of stress susceptibility.

## INTRODUCTION

Extensive epidemiological evidence links stressful life experiences with depression, a chronic, episodic condition characterized by low mood, anhedonia and diminished motivation among other symptoms (Billings et al., 1983). The physiological systems that maintain body functions to meet external demands can become dysregulated by chronic stress, increasing ‘allostatic load’ (McEwen & Stellar, 1993). This disrupts neuroendocrine feedback mechanisms and precipitates neural adaptations across various brain regions that enable cognitive, appetitive and emotional processes (reviewed in: Parekh et al., 2022). These adaptations include structural and functional changes to cellular components critical for information processing and neuronal output. While many depressed patients exhibit reduced hedonic capacity or decreased “liking” of rewards, amotivation is distinct and characterized by reduced capacity to pursue rewards or decreased “wanting” (Der-Avakian & Markou, 2012; Nusslock & Alloy, 2017; Treadway & Zald, 2011). Amotivation is particularly relevant in depression and involves the complex interaction of limbic and cognitive processes (Cooper et al., 2018). Existing preclinical assays, such as the sucrose preference test, tend to focus on consummatory anhedonia and do not model anticipatory or motivational domains of dysfunction seen in clinical conditions (Dichter et al., 2010). Therefore, it is important to evaluate a variety of behavioral outcome measures to assess the impact of stress on motivation. To this aim, an individual’s willingness to engage in effortful goal-directed behavior is an important consideration. Reward pursuit despite effort requirements is a core feature of adaptive reward-seeking (Le Heron et al., 2018). Patients who suffer from amotivation experience deficits in such cost-benefit analyses, affecting engagement, regardless of how pleasantly they may perceive rewards once consumed (Salamone et al., 2016; Treadway et al., 2012; X. Yang et al., 2014). Chronic stressors impair reward salience and effortful responding in rodent operant tasks, however, the underlying neural and neurobiological mechanisms are incompletely understood (Bryce & Floresco, 2016; Dieterich et al., 2021; Kúkel’ová et al., 2018; Xeni et al., 2024).

The anterior cingulate cortex (ACC) integrates a variety of signals pertaining to reward values, reinforcement history and expected effort expenditure to drive reward seeking or disengagement under changing contingencies (Paus, 2001; Kennerley et al., 2006; Rushworth & Behrens, 2008; Klein-Flügge et al., 2016). It is ideally positioned to perform these computations given its connections with the ventral tegmental area (VTA), nucleus accumbens (NAc), amygdala, hippocampus and neighboring prefrontal regions (Rudebeck et al., 2006)(André et al., 2019). Dorsal ACC lesions in neurological patients can cause abulia and avolition (Devinsky et al., 1995; Grunsfeld & Login, 2006) while, interestingly, the subgenual ACC is hyperactive in MDD patients and is a target for putatively inhibitory neuromodulatory and drug therapies, implying the human ACC plays a nuanced and subregion-specific role in the etiology of motivational deficits and depression (Mayberg et al., 2005; Alexander et al., 2019, 2021). In rodents, ACC inactivation specifically affects the amount of effort invested for rewards, biasing animals towards suboptimal low effort/low reward responses, despite intact discrimination (Schweimer & Hauber, 2005; Floresco & Ghods-Sharifi, 2007; Floresco et al., 2008). We recently established the causal role of an ACC-NAc pathway in integrating learned effort and reward information to drive motivated responding and demonstrated that chronic neuroendocrine disruption decreases high-effort/high-reward choices and circuit activity (Fetcho et al., 2024). Interestingly, circuit dysfunction scaled with the severity of the behavioral deficit, suggesting stress-sensitive ACC projections may contribute to individual variability in stress response, though potential contributing molecular mechanisms were not explored.

The anterior cingulate cortex (ACC) integrates a variety of signals pertaining to reward values, reinforcement history and expected effort expenditure to drive reward seeking or disengagement under changing contingencies (Paus, 2001; Kennerley et al., 2006; Rushworth & Behrens, 2008; Klein-Flügge et al., 2016). It is ideally positioned to perform these computations given its connections with the ventral tegmental area (VTA), nucleus accumbens (NAc), amygdala, hippocampus and neighboring prefrontal regions (Rudebeck et al., 2006)(André et al., 2019). Dorsal ACC lesions in neurological patients can cause abulia and avolition (Devinsky et al., 1995; Grunsfeld & Login, 2006) while, interestingly, the subgenual ACC is hyperactive in MDD patients and is a target for putatively inhibitory neuromodulatory and drug therapies, implying the human ACC plays a nuanced and subregion-specific role in the etiology of motivational deficits and depression (Mayberg et al., 2005; Alexander et al., 2019, 2021). In rodents, ACC inactivation specifically affects the amount of effort invested for rewards, biasing animals towards suboptimal low effort/low reward responses, despite intact discrimination (Schweimer & Hauber, 2005; Floresco & Ghods-Sharifi, 2007; Floresco et al., 2008). We recently established the causal role of an ACC-NAc pathway in integrating learned effort and reward information to drive motivated responding and demonstrated that chronic neuroendocrine disruption decreases high-effort/high-reward choices and circuit activity (Fetcho et al., 2024). Interestingly, circuit dysfunction scaled with the severity of the behavioral deficit, suggesting stress-sensitive ACC projections may contribute to individual variability in stress response, though potential contributing molecular mechanisms were not explored.

To explore this, we used a novel conditioned reward seeking task in which animals’ motivation to pursue rewards in the face of an effort cost can be determined by measuring both anticipatory and consummatory responses. Unsupervised clustering revealed that exposure to chronic psychosocial stress reduced anticipatory responding on trials requiring low and high effort in a subset of animals termed ‘susceptible’, while in others this measure was largely unaffected by stress exposure (‘resilient’). Spatial transcriptomics analysis of the ACC from control and stressed animals revealed divergent cell-type specific transcriptional signatures enriched for synapse-related gene sets in susceptible and resilient mice. Lastly, spatially restrained cell-cell communication analysis identified altered astrocytic and inhibitory signaling in the stress-susceptible animals as putative sources of altered ACC function upstream of behavioral changes.

## METHODS

### Animals

Male and female C57Bl/6 mice (Jackson Labs) were aged 6-7 weeks at the time of surgery. Retired breeder male CD1 mice (Charles River) aged 3-4 months were used for chronic non-discriminate social defeat stress (CNSDS). All animals were maintained on a 12:12 reverse light/dark cycle (lights off at 9:00AM) with free access to food and water except during periods of water restriction (see ‘Head-restrained Effortful Reinforcement Task’ section). Experimental procedures were approved by the Weill Cornell Medicine Institutional Animal Care and Use Committee and conducted in accordance with NIH guidelines for the care and use of laboratory animals.

### Headplate surgery

Mice were briefly anesthetized by inhalation of isoflurane mixed with oxygen (induction, 5%; maintenance, 1-2%). Dexamethasone (1mg/kg; i.p.) and meloxicam (1mg/kg; i.p.) were administered prior to surgery to reduce brain swelling and as a prophylactic analgesic, respectively. Once placed on a stereotaxic frame (Kopf Instruments), sterile lubricating ointment (Puralube) was applied to the eyes, and body temperature was maintained using a microwaveable heating pad (Snugglesafe). Scalp fur was cleaned and trimmed, and skull was exposed with a midline scalp incision. Bupivicaine (0.05mL; 5mg/mL) was applied to the skull surface as a secondary analgesic. A custom lightweight titanium headplate was then secured to the skull with Metabond (Parkell, Inc.) for head-restrained behavioral training.

### Head-restrained Effortful Reinforcement Task

Mice were water restricted within 90% of pre-restriction body weight and handled daily for 1 week then habituated to head-fixation on a custom-built Arduino-controlled behavioral setup. During lickspout habituation (1-2 days), 3-4uL water drops were freely dispensed every 2-seconds until animals reached criterion licking (>500 licks/session). Once habituated, mice were trained to discriminate two auditory tones of differing frequencies (CS- and CS+; 6kHz, 13kHz - counterbalanced) predicting omission or presentation of a water drop reward. The reward cue was presented for 2-seconds followed by a trace interval (TI) period of 2-seconds and reward delivery or omission. A variable inter-trial-interval (ITI; 8,10 or 12-seconds) between trials was used to prevent habitual responding. Once mice displayed adequate reward cue discrimination, using d-prime (d’) as the discrimination index (Z(hit rate) - Z(false alarm rate)), the full-task training protocol was initiated. In addition to reward cues, a tactile whisker brush cue of different stroke patterns (pulsed or sweeping pattern; counterbalanced) was introduced and co-terminated with the auditory tone. This signaled the effort requirement of the trial. On low-effort trials, the lickspout remained parked while on high-effort trials, the lickspout rotated downward at an angle from the parked position (2-4 degrees) which was empirically determined for each individual animal to be at a distance requiring full tongue extension for access. Sessions consisted of 150 pseudo-randomly presented trials. Timestamps of individual lick responses were continuously recorded, and trial data was parsed using custom Python scripts. Anticipatory and consummatory responses were quantified by subtracting the number of licks made during the TI or reward omission/delivery period (5-seconds) from the last 2-seconds of the ITI, respectively. Data were processed using custom MATLAB code and imported to GraphPad Prism for plotting and statistical analysis. The change in anticipatory licking after chronic stress (described below) was used as input into unsupervised K-means clustering (k=2; performed in R) to classify animals as susceptible or resilient. The total number of clusters was determined using an elbow-plot of the total within-cluster sum of squares at values of k ranging from 1 to 5.

### Chronic non-discriminatory social defeat stress (CNSDS)

Animals of both sexes were subjected to a multi-day protocol to induce psychosocial stress as previously described (Yohn et al., 2019). Briefly, test mice (one male and one female C57Bl/6) were exposed to daily encounters (5-10 min) with a larger aggressor mouse (male CD1) in the latter’s home cage (measuring 27 x 48 x 15cm with woodchip bedding). This reliably elicits an antagonistic interaction characterized by social dominance behaviors by the aggressor mouse and submissive and coping behaviors by the test mice. We documented that a larger proportion of attacks are directed towards male test mice (82.7%) compared with females (17.3%) (**Suppl. Figure 1A**) in accord with results described by Yohn and colleagues though we did not find a significant difference in attack latency (**Suppl. Figure 1B**; Unpaired t test; p=0.2881). After each encounter, a test mouse and aggressor remain housed together, separated by a perforated Plexiglas divider, in the aggressor’s home cage for 24 hours to prolong a sensory stress experience. Male and female test mice alternate overnight housing with their paired aggressor versus another CD1. They are then moved to the home cage of a new aggressor the following day, and this procedure is repeated daily for 10 days to prevent habituation. Defeat sessions were video recorded and scoring of attacks and attack latency were performed by an experimenter blinded to animal IDs and groups. Control animals remained in their home cages and were handled daily.

### Open Field Social Interaction (OFSI)

One day following the end of the social defeat or stress control protocol, mice were tested for social interaction behaviors using an established protocol (Golden et al., 2011). Briefly, mice were habituated to the behavior room for at least 30 minutes under dim lighting conditions. An opaque Plexiglas arena measuring 40cm x 40cm x 40cm was used for the assay and during the first phase, a clear Plexiglas enclosure measuring 10cm x 6cm x 40cm with slits all around was placed centrally against one wall. An experimental mouse was introduced into the arena and allowed to explore for 150s after which it was removed into the home cage for 30 seconds and the enclosure was replaced with one containing a novel aggressive CD1 mouse. The experimental mouse was reintroduced to the arena for 150s. Activity patterns in the ‘interaction zone’ around the enclosure and rear ‘corner zones’ were video recorded and tracked automatically using Ethovision XT 11.5 software (Noldus). A social interaction (SI) score was calculated as the time spent interacting with the CD1 aggressor in the interaction zone in the second phase versus time spent interacting with the empty enclosure during the first phase.

### Elevated Plus Maze (EPM)

To assess anxiety-like and exploratory behaviors, animals were tested in a custom-built plus-shaped maze consisting of four white Plexiglas arms measuring 29cm x 6cm. Two opposing arms included opaque walls measuring 14cm in height (closed arms). The maze was elevated 70cm above the ground. Prior to testing, mice were habituated to the behavior room for at least 30min under dim lighting conditions and the center of the maze was illuminated to ∼10 Lux. To begin each trial, a mouse was placed at the center facing an open arm and allowed to explore the entire maze for 10min. Mice were video recorded from above and their movements tracked using Ethovision XT 11.5 software (Noldus). The amount of time spent in the open arms and well as percentage of open arm entries normalized to total crossings (all four paws within an arm) were calculated.

### Tissue preparation

Animals were rapidly euthanized by cervical dislocation followed by decapitation. Brains were removed into cold artificial cerebral spinal fluid (aCSF), rinsed and crudely sectioned in a block (F.S.T.). The ACC was micro-dissected and placed into 4% paraformaldehyde for fixation. A standard protocol was followed to prepare formalin fixed paraffin embedded (FFPE) tissue from mouse brain (Smart et al., 2023) and ACC pieces were arranged in an array (n = 9, 3/group, 2 males and 1 female) and serially sectioned on a microtome. Two slices containing the ACC array were mounted on 10X Visium v1 slides (Transcriptome Probe Set v1.0) and processed according to the manufacturer’s protocol. Slides were imaged under an epifluorescence microscope (Leica Microsystems) to identify the fiducial boundaries of the capture sites using a TRITC filter.

### Spatial RNA sequencing and analysis

Library preparation and sequencing was performed in collaboration with Weill Cornell Medicine’s Epigenomics Core. Sequencing data were aligned to the 10x probe set using SpaceRanger (v2.0.0) and fluorescent images of the tissue were manually aligned to the fiduciary boundaries. The set of spots corresponding to the ACC of each replicate in the tissue array was manually obtained using the Loupe browser (10X Genomics). Alignment of biological replicates into a single coordinate space (for visualizations), as well as clustering of spot-level data, was performed using SPIRAL (Guo et al., 2023). Spot-level data analysis was performed using the spot x gene matrix obtained from SpaceRanger and processing in Seurat (Hao et al., 2024). Cell-level data analysis was performed by deconvolution of the spot x gene matrix using Cell2Location (Kleshchevnikov et al., 2022), with a large single-cell RNA-seq dataset (n = 59,218 cells) of the ACC used as the reference (Yao et al., 2021). For the Cell2Location user-defined parameters, we selected detection_alpha=200 and N_cells_per_location=8. The cell x gene matrix containing the predicted cell-level expression of cells within spots was further processed using Seurat. Processing in Seurat of both spot- and cell-level data included data normalization using SCTransform, and dimensionality reduction with PCA and UMAP. Differential expression analysis between pairs of the three groups (control, susceptible, and resilient) was performed by using the FindMarkers function in Seurat within each cell-type or spot cluster identity, and significance was defined as padj < 0.05. Visualization of the overlaps between cell type- and group comparison-specific DEGs was performed using UpSet plots (Lex et al., 2014), and heatmap visualization and analysis was performed using ComplexHeatmap (Gu et al., 2016). Enrichment analysis of DEG lists was performed using gProfiler (Kolberg et al., 2023). Cell-cell communication analysis was performed using CellChat v2 (Jin et al., 2024), with spatial coordinate information used as a restraint for secreted signaling (250μm) and contact signaling (55μm, the diameter of a Visium spot). Downstream analysis of CellChat results was performed in Seurat by sub-setting groups of cells based on their cell type, their expression of ligand-receptor molecules, and their proximity to other cell types expressing the corresponding ligand-receptor and then performing differential expression between groups using the FindMarkers function.

### Statistical analysis

Statistical analyses were conducted in GraphPad Prism Version 10.3.1 and in R (https://www.r-project.org/). Behavioral analyses used Unpaired and Paired t tests, one-way ANOVA and 2-way ANOVA where appropriate. For differential expression analyses, the FindMarkers function in Seurat was used with default settings (Wilcoxon Rank Sum Test with the Benjamini-Hochberg correction for multiple comparisons). For cell-cell communication analyses, pathways with significant changes in predicted signaling between selected cell types (i.e. **Figure 5C**, **Figure 6A, Suppl. Figure 5D, Suppl. Figure 5H**) were determined using the do.stat=TRUE option in the rankNet function in CellChat, which compares signaling probabilities between groups using the Wilcoxon Rank Sum Test with the Benjamini-Hochberg correction for multiple comparisons.

### Data availability

10X Visium sequencing data from this study will be made publicly available upon publication.

## RESULTS

### Chronic psychosocial stress differentially affects motivated responding during effortful reward seeking

We aimed to elucidate cell type-specific transcriptional effects of chronic stress within the ACC. Stress effects were evaluated with our novel conditioned reward-seeking task, which allowed us to measure anticipatory and consummatory responses for a water reinforcer in water-restricted mice of both sexes, as well as sensitivity to expected reward availability and effort requirements. Mice were habituated and trained in stages (**Figure 1A**). In the full task, animals are presented with cued reinforced (CS+) and non-reinforced (CS-) trials in which the water-dispensing lick-spout is positioned closer to the mouth during “low effort (LE)” trials or at a greater distance on “high effort (HE)” trials (**Figure 1B – left**). Auditory tones of differing frequency serve as reward cues and whisker brushes of varying sweep patterns signal effort condition (counterbalanced across subjects). Cues co-terminate and are followed by a 2-second trace interval (TI), reward delivery/omission period (5s) and variable duration inter-trial interval (ITI) (**Figure 1B - right**). Mice reliably learned to discriminate reward cues from early to late training sessions (**Figure 1C**). Moreover, animals modulated their anticipatory lick responses during the trace interval period depending on expected reward availability and level of difficulty (**Figure 1D,E**). They displayed a similar and expected pattern of consummatory licking on each trial type (**Figure 1F**).

**Figure 1.**
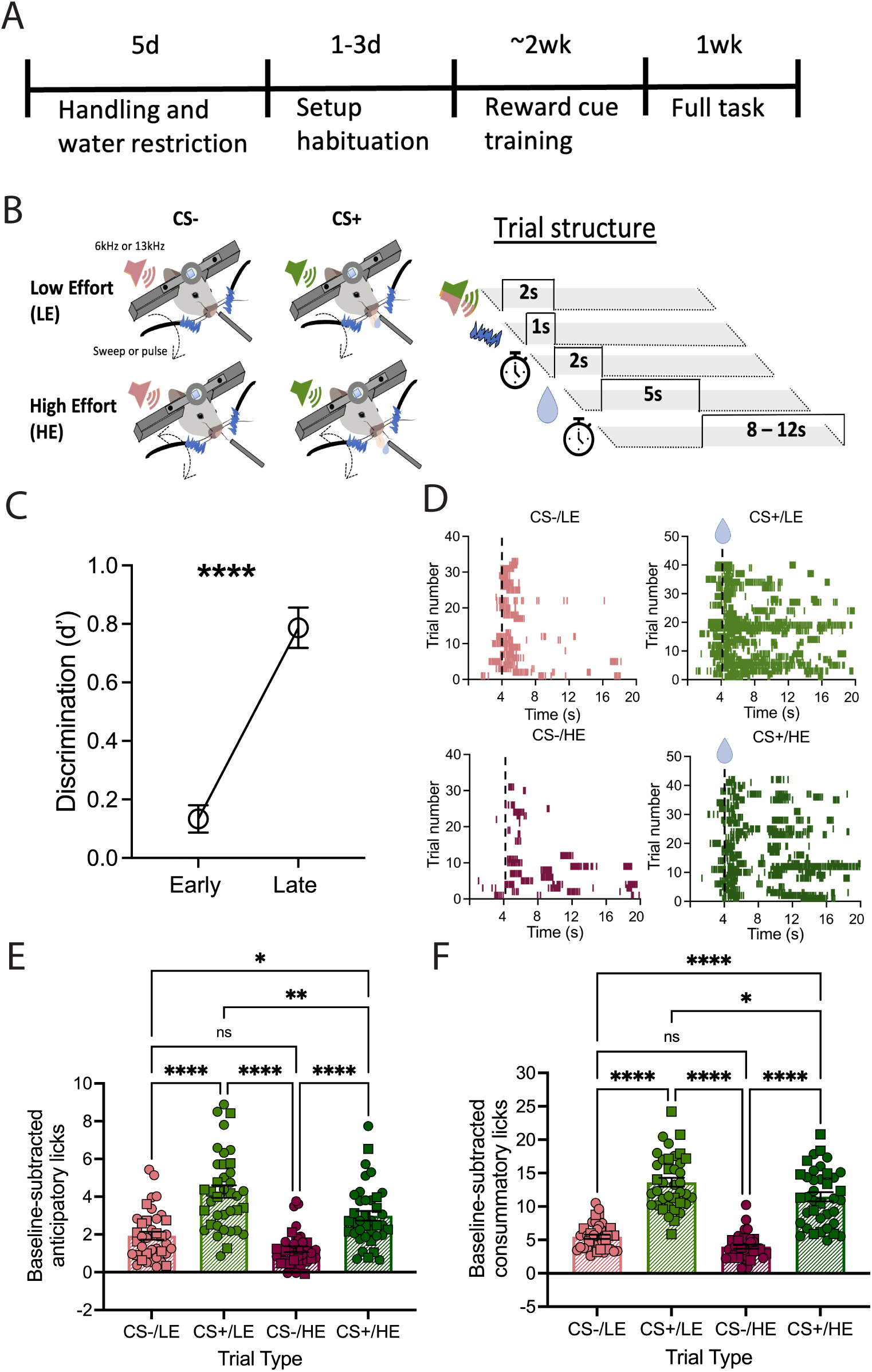
A head-restrained effortful reinforcement task. **A**) Schematic overview of the training schedule. **B**) Four trial types presented in the task (right) include CS-/LE (unrewarded/low-effort), CS+/LE (rewarded/low-effort), CS-/HE (unrewarded/high-effort) and CS+/HE (rewarded/high-effort) and the trial structure (right) with epochs (in seconds) including cue presentation, trace interval delay period (anticipatory), reward delivery or omission period (consummatory) and variable duration inter-trial interval (ITI). **C**) Cue learning as measured by d’ discrimination from early to late training sessions (n=38 mice; 3 sessions each; Paired t test p<0.0001). **D**) Representative raster plots depicting licks made during each of the trial types in a single session. **E**) Anticipatory licks made in response to each trial type averaged across 5 sessions (n=38 mice; 13 females (square symbols) one-way ANOVA F(3,148)=28.092; p<0.0001)). **F**) Consummatory licks made in response to each trial type averaged across 5 sessions (one-way ANOVA F(3,148)=77.73; p<0.0001). *p<0.05,**p<0.01; ****p<0.0001.

We next employed 10 days of chronic non-discriminatory social defeat stress (**Figure 2A**; **Suppl. Figure 1**) with a subset of animals remaining unexposed (Control group). While CNSDS did not significantly affect social interaction as has been reported (Yohn et al., 2019; **Suppl. Figure 1C**), we observed a trend level effect on open arm entries in the elevated plus maze (EPM) and significant reduction in open arm time in stressed animals (**Suppl. Figure 1D-E**). In this task, we found that CNSDS reduced anticipatory responding on trials in which animals expended either low or high levels of effort for the reinforcer (CS+/LE and CS+/HE trials) while the reward-seeking behavior of control animals remained unchanged over time (**Figure 2B-E)**. Notably, while we observed a decrease in anticipatory licking in stressed mice **(Figure 2F),** the effect of stress was continuous and some stressed mice exhibited anticipatory licking similar to controls. To group mice along this continuum we performed unsupervised K-means clustering of anticipatory licking during high-effort, rewarded trials (CS+/HE). Plotting the Within-Cluster Sum of Squares (WCSS) against the number of clusters (k) revealed an elbow at k=2 (**Figure 2G**), and K-means clustering with k=2 identified a cutoff at which we could split the stressed mice into two groups with distinct changes in anticipatory licking after stress (**Figure 2H**). For downstream analyses we labeled animals belonging to cluster 1, whose anticipatory licking was relatively unaffected by stress, ‘resilient’, and animals belonging to cluster 2 as ‘susceptible’. Interestingly, we found that a relative change in reward seeking in the conditioned task did not correlate with social interaction or anxiety-related behaviors in stressed mice (**Suppl. Figure 2**). We then asked whether we could identify underlying gene-regulatory network-level differences corresponding to differential post-stress behavioral profiles.

**Figure 2.**
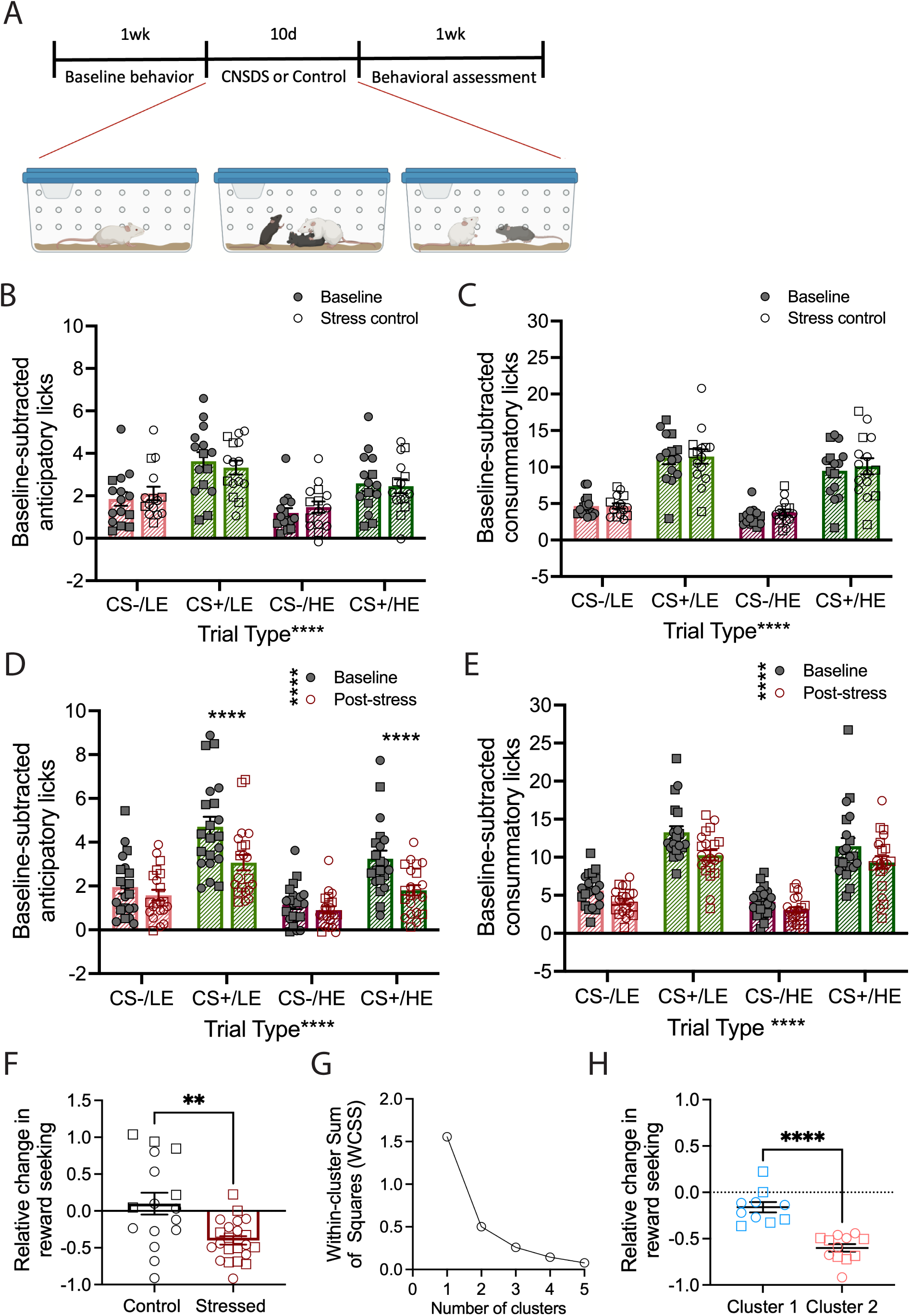
Effects of chronic psychosocial stress on effortful reinforcement behavior. **A**) Schematic overview of the timeline of behavioral testing and the chronic nondiscriminatory social defeat stress (CNSDS) protocol. **B**) Average anticipatory licks made for each trial type at baseline and after 10 days of normal housing with gentle handling for control animals (2-way ANOVA; trial type: F(3,56)=10.59; p<0.0001; treatment: F(1,56)=0.01796; p=0.8939; n=15 control, n=22 stressed). **C**) Average consummatory licks across trial types for control animals (2-way ANOVA; trial type: F(3,56)=34.29; p<0.0001; treatment: F(1,56)=1.008; p=0.3198). **D**) Average anticipatory licking in CNSDS exposed animals compared with baseline behavior (2-way ANOVA; trial type: F(3,80)=19.46; p<0.0001; treatment: F(1,80)=54.02; p<0.0001); interaction: F(3,80)=8.142; p<0.0001). **E**) Average consummatory licking in CNSDS exposed animals compared with baseline behavior (2-way ANOVA; trial type: F(3,80)=56.29; p<0.0001; treatment F(1,80)=23.48; p<0.0001; interaction: F(3,80)=1.183; p=0.3214). **F**) Relative change in reward seeking (anticipatory licking on low- and high-effort rewarded trials) for control and CNSDS animals (Unpaired t test; p=0.0052; n=37 mice). **G**) Elbow plot from K-means clustering. **H**) Distinct clusters identified by K-means clustering on the relative change in reward seeking in stressed mice (Unpaired t test; p<0.0001). **p<0.01; ****p<0.0001. Panel A created with BioRender.com

### Spatial RNA-sequencing of ACC reveals distinct sets of differentially expressed genes correlated with motivational impairment

We hypothesized that resilience to stress-induced motivational deficits is marked by a unique transcriptional signature within the ACC. We employed a spatial RNA-sequencing platform (10X Visium) to test this (**Figure 3A**). This approach assesses the transcriptional profile of hexagonal segments of the tissue sections (55um diameter), called spots, which typically contain an admixture of multiple cells. Analysis of spot-level data revealed four clusters of spots which corresponded to cortical layers (**Suppl. Figure 3A**) and were evenly distributed across Visium slides and groups (**Suppl. Figure 3B**). Cluster 1, corresponding to the midline of the brain and layer 1 of the cortex, displayed marker gene expression consistent with primarily astrocyte and endothelial cell identity, while clusters 2, 3, and 4 had marker gene expression consistent with cortical layer 2/3, layer 5, and layer 6 excitatory neurons, respectively (**Suppl. Figure 3C-D**). To identify gene expression changes associated with stress susceptibility and exposure, we first performed differential expression analysis within spot clusters between control and susceptible (C-S), control and resilient (C-R) and susceptible and resilient (S-R) groups, revealing on average 10 differentially expressed genes (DEGs) for each cluster and comparison (**Suppl. Figure 3E**). These DEGs included Sgk1, a well-established stress responsive gene (Licznerski et al., 2015), which is downregulated after chronic stress in both resilient and susceptible animals within cluster 3 (layer 5; **Suppl. Figure 3F**). Other examples include Gfap, a maker of astrocyte identity as well as astrocyte reactivity (Z. Yang & Wang, 2015), which was downregulated in resilient animals compared to the other two groups in clusters 2 & 3 (layers 2/3 and 5, respectively; **Suppl. Figure 3F**), as well as Nr4a3, an immediate early gene and transcriptional regulator (Campos-Melo et al., 2013), which was downregulated in susceptible animals compared to controls in cluster 4 (layer 6; **Suppl. Figure 3F**). In sum, ACC tissue spots clustered into cortical layers and exhibited modest changes in gene expression which included known stress-sensitive genes.

**Figure 3.**
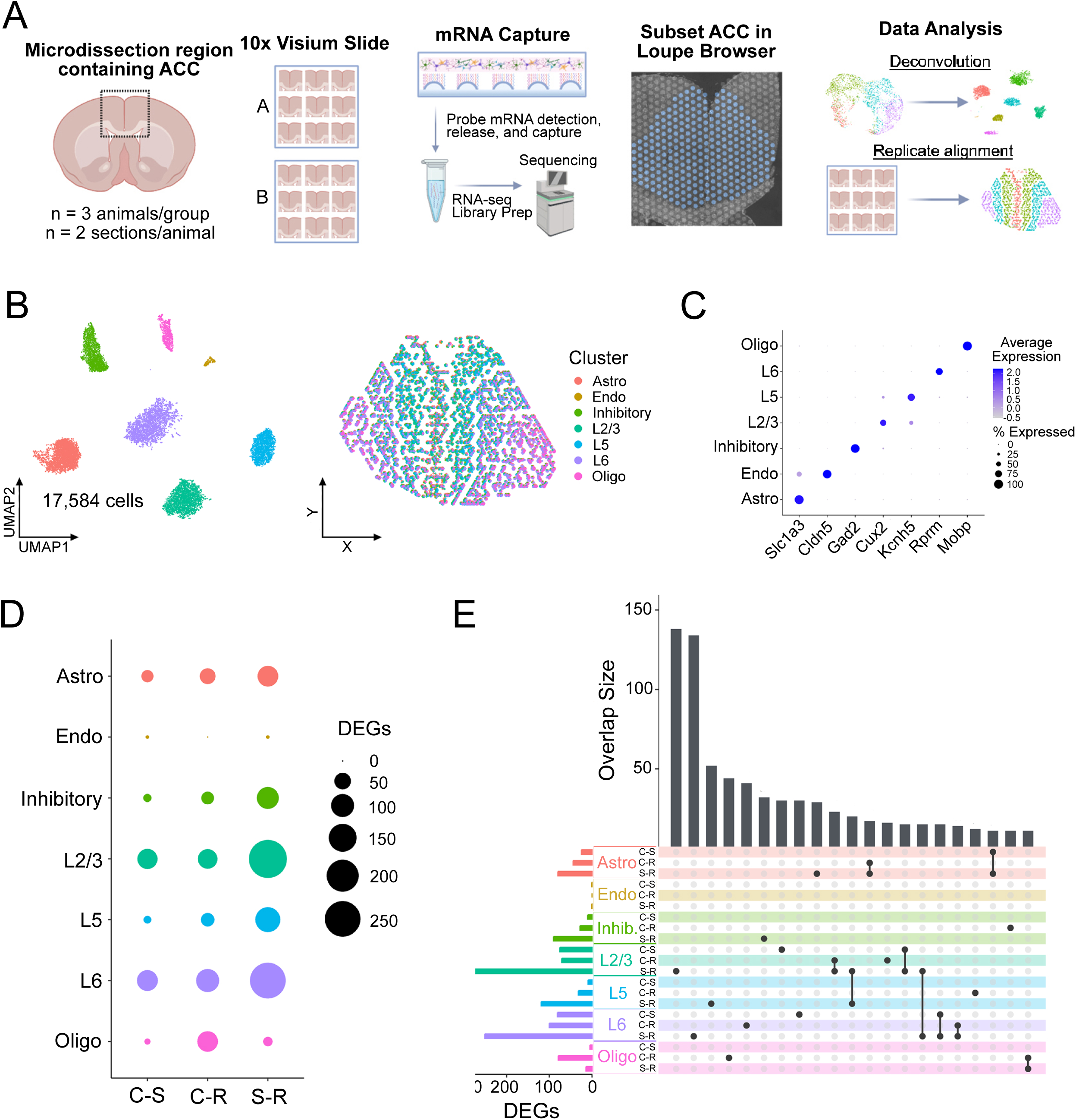
Cell type-specific transcriptional effects of chronic stress in the ACC. **A**) A scheme depicts the study design for the spatial transcriptomics experiment. **B**) Deconvoluted cells detected in the Visium dataset shown on a UMAP plot (left) and a consensus spatial plot (right). **C**) A marker gene dotplot depicts 7 genes whose expression is largely confined to one of the 7 identified cell clusters. **D**) A dotplot depicts the number of DEGs identified in each cell cluster for each of the three paired group comparisons (C-S: control vs. susceptible; C-R: control vs. resilient; S-R: susceptible vs. resilient). **E**) An UpSet plot shows the top 21 overlapping DEG lists (ordered by size) between each comparison within each cell cluster. A single point denotes unique DEGs and two or more points connected by a line denotes genes shared between those comparisons. Panel A created with BioRender.com.

In order to measure group-level expression changes while accounting for the heterogeneity of cell types within each tissue spot, we performed deconvolution using Cell2Location (Kleshchevnikov et al., 2022) with a large ACC scRNA-seq dataset as a reference (**Suppl. Figure 4**) (Yao et al., 2021). Deconvolution successfully separated astrocytes, endothelial cells, inhibitory neurons, oligodendrocytes, and layer-specific pyramidal cells from the spot-level data (**Figure 3B**), and each deconvolved cell type exhibited high expression of the expected marker genes (**Figure 3C**). Importantly, cell deconvolution allowed us to detect a greater number of DEGs compared to the spot-level analysis (**Figure 3D**). An initial overlap of cluster-and comparison-specific DEG lists revealed that the majority of these DEGs were specific to both cluster and group comparison. However, we did observe overlaps both across group comparisons in the same cluster as well as within group comparisons across clusters (**Figure 3E**). Together, these results demonstrate that susceptibility and resilience to stress-induced motivational impairments are distinguished by unique patterns of cell-type specific differentially expressed genes within the ACC.

To identify broad patterns of stress-related differential gene expression, we focused on classifying DEGs that overlap across group comparisons within each cell cluster. We therefore constructed heatmaps for each cell type composed of genes identified as DEGs in at least 2 out of 3 group comparisons (**Figure 4A**). This analysis revealed three distinct patterns of overlapping DEGs: 1) genes altered concordantly in both resilient and control groups compared to susceptible animals, which we call ‘Susceptibility genes’, 2) genes altered concordantly in both susceptible and control groups compared to resilient animals, which we call ‘Resilience genes’, and 3) genes altered concordantly in both susceptible and resilient groups compared to control mice, which we call ‘Stress genes’ (**Figure 4A**). These categories of genes exhibited distinct gene ontology enrichment, with Resilience genes enriched for terms related to synapses and glutamatergic function, Susceptibility genes enriched for vesicle regulation terms, and Stress genes enriched for biological processes including hormone response, cognition, and behavior (**Figure 4B**). Representative examples of Resilience genes include Adcyap1 in L2/3 cells, encoding the PACAP neuropeptide that is critically involved in the stress response (Martelle et al., 2021), and Ppp3r1 in astrocytes, encoding the calcium-dependent phosphatase calcineurin, plays a role in neuronal resilience via anti-inflammatory actions in astrocytes (**Figure 4C**, **Suppl. Figure 5A**) (Fernandez et al., 2007). Susceptibility genes include Lmo4 in inhibitory neurons, encoding a transcriptional regulator implicated in reward learning and stress-induced anxiety-related behaviors (Maiya et al., 2015; Qin et al., 2015), and Lin7a in L6 neurons, encoding a scaffold protein involved in synaptic signaling and neurite growth (**Figure 4D**, **Suppl. Figure 5B**) (Crespi et al., 2012; Olsen et al., 2006). Stress genes include Sgk1 in L6 neurons, which we also identified in the spot-level data (**Suppl. Figure 2F**), and Ptgds in astrocytes, encoding a prostaglandin synthase that has been identified as a potential marker for reactive astrocytes (**Figure 4E**, **Suppl. Figure 5C**) (Matusova et al., 2023). Together, our differential expression analysis in the ACC is in line with previous reports of the transcriptional effects of stress in frontal cortical regions, which similarly find a majority of DEGs in excitatory neurons (Hing et al., 2024; Z. Yang & Wang, 2015), altered expression of synapse-related gene sets (Licznerski et al., 2015; Kwon et al., 2022), and a role for astrocytes in mediating stress susceptibility (Holt et al., 2024).

**Figure 4.**
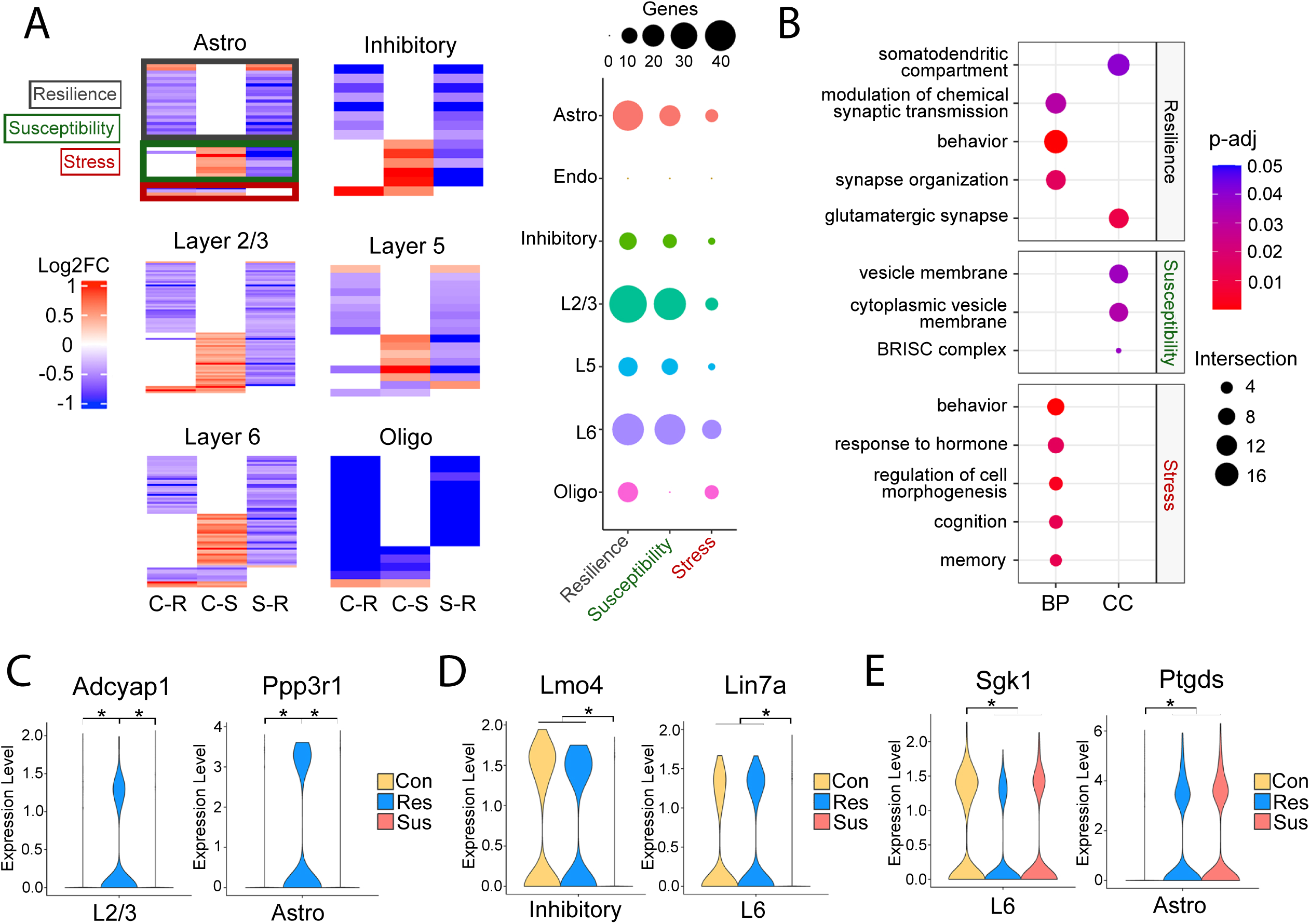
Chronic stress produces divergent transcriptional signatures in susceptible and resilient animals. **A**) Heatmaps show the log_2_ fold-change (Log2FC) of DEGs that are shared across group comparisons within each cell cluster (left; C-S: control vs. susceptible; C-R: control vs. resilient; S-R: susceptible vs. resilient). From these heatmaps we identified 3 general patterns: Resilience genes change in the same direction in the C-R and S-R comparison (black box), Susceptibility genes change in the opposite direction in the C-S and S-R comparisons (green box), and Stress genes change in the same direction in the C-R and C-S comparison (red box; patterns are boxed in the astrocyte heatmap, top-left, as an example). A dotplot on the right shows the number of genes in each of these three categories identified in each cell type. **B**) An enrichment dotplot shows significantly enriched gene ontology terms in the Biological Process (BP) and Cellular Compartment (CC) categories in each of the three patterns of overlapping DEGs. Violin plots show the expression of the Resilience genes *Adcyap1* in L2/3 cells and *Ppp3r1* in astrocytes (**C**), Susceptibility genes *Lmo4* in inhibitory neurons and *Lin7a* in L6 cells (**D**), and Stress genes *Sgk1* in L6 cells and *Ptgds* in astrocytes (**E**). *p_adj_ < 0.05, Wilcoxon Rank Sum test, Benjamini-Hochberg correction).

### The strength of inhibitory signaling within ACC microcircuitry correlates with resilience to stress-induced motivational impairment

Next, we tested whether the additional spatial dimension present in our dataset could be leveraged to identify cell-cell communication patterns upstream of the observed changes in gene expression. To this end, we assessed cell-cell communication probability using the CellChat package (Jin et al., 2024), with the spatial proximity between cells used as a restraint for inferred signaling. This analysis revealed similar overall interaction strength in susceptible compared to control animals (**Suppl. Figure 6A-C**), while interaction strength was elevated in resilient animals compared to both control (**Suppl. Figure 6E**) and susceptible animals (**Figure 5A**). This was driven by inhibitory to excitatory neuron interactions in both cases (**Figure 5B**, **Suppl. Figure 6F-G**), which included GABA-B signaling as a pathway with significantly elevated interaction strength in resilient animals compared to control (**Suppl. Figure 6H**) and susceptible animals (**Figure 5C-D**). In addition to GABA-B, several neuropeptide signaling pathways exhibited elevated strength in the ACC neurons of resilient compared to susceptible animals (**Figure 5C**). Of these, we focused further on Crh, which encodes corticotropin releasing hormone (CRH), due to its well-established role in the physiological stress response and effects on cognitive function (Hupalo, Bryce, et al., 2019). We also explored opioid signaling, as recent reports implicate various types of cortical opioid signaling with the effects of stress and antidepressant treatments (Mallimo & Kusnecov, 2013; Williams et al., 2018, Y.-J. Wang et al., 2023; Jiang et al., 2024, Munguba et al., 2026). Interestingly, the signaling strength of these pathways were not significantly altered in the control vs. susceptible comparison (**Suppl. Figure 6D**) or the control vs. resilient comparison (**Suppl. Figure 6H**), implying Crh and opioid signaling in the ACC may therefore play a specific role in mediating resilience in stress-exposed animals.

**Figure 5.**
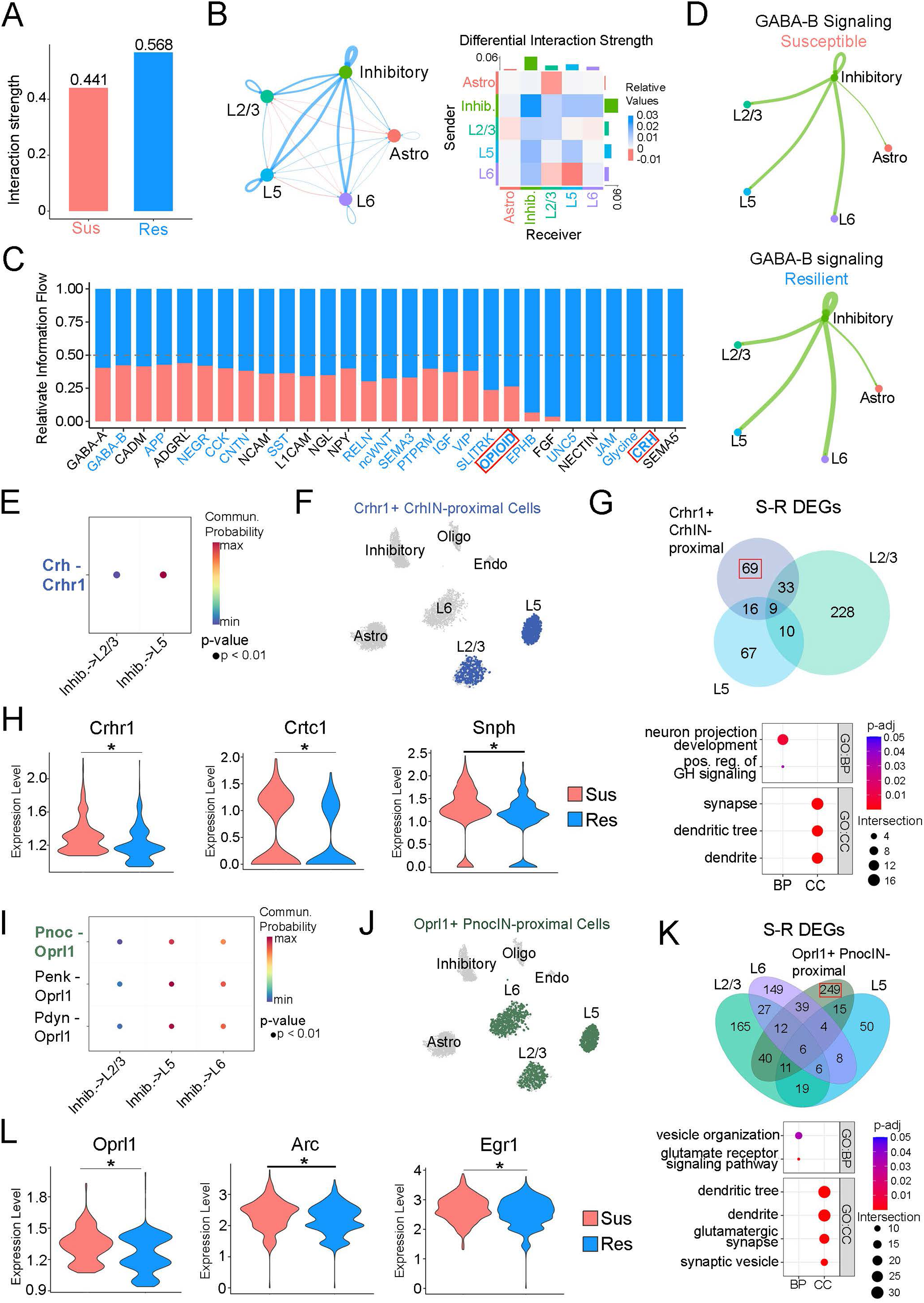
Stress resilience is marked by enhanced neuropeptide signaling from inhibitory to excitatory neurons. **A**) A barplot shows the difference in overall interaction strength determined by CellChat between the susceptible and resilient groups. **B**) A circle plot (left) and heatmap (right) show the differential interaction strength between specific cell clusters, with red and blue representing interactions with increased probability in the susceptible condition (compared to resilient) or vice versa, respectively. Edge width and color intensity represent the magnitude of difference between groups in the circle plot and heatmap, respectively. **C**) A stacked bar plot depicts the difference in relative signaling strength (information flow) between the susceptible and resilient conditions for specific signaling pathways with inhibitory neurons as the source and excitatory neurons as the targets (blue x-axis labels denote a statistically significant increase in signaling strength in the resilient compared to susceptible condition). **D**) Circle plots show the predicted strength of GABA-B signaling (significantly altered in **C**) between cell types in the susceptible (top) and resilient (bottom) conditions. **E**) A dotplot demonstrates that the predicted inhibitory to excitatory Crh signaling (boxed in **C**) consists of the Crh-Crhr1 ligand-receptor (LR) pair and occurs between inhibitory neurons and L2/3 and L5 excitatory neurons (data from the resilient group shown as an example). **F**) A UMAP plot highlighting L2/3 and L5 cells that both 1) express *Crhr1* and 2) are within close proximity (250μm) of a *Crh*+ inhibitory neuron (CrhIN). **G**) A Venn diagram (top) depicts the overlap of susceptible vs. resilient (S-R) DEGs in the cells highlighted in **F** with the general populations of L2/3 and L5 neurons, along with a dot plot (bottom) showing the enrichment of gene ontology Biological Process (BP) and Cellular Component (CC) terms in the 69 DEGs specific to Crh receiver cells (boxed above). **H**) Examples of genes specifically altered in the S-R comparison of Crh receiver cells include *Crhr1*, *Crtc1*, and *Snph*. **I**) A dotplot demonstrates that the predicted inhibitory to excitatory opioid signaling (boxed in **C**) includes Pnoc-Oprl1 as an LR pair and includes all 3 groups of excitatory neurons as signaling targets for Pnoc signaling (data from the resilient group shown as an example). **J**) A UMAP plot highlighting excitatory cells that both 1) express *Oprl1* and 2) are within close proximity (250μm) of a *Pnoc*+ inhibitory neuron (PnocIN). **K**) A Venn diagram (top) depicting the overlap of S-R DEGs in the cells highlighted in **J** with the general populations of layer-specific excitatory neurons, along with a dot plot (bottom) showing the enrichment of BP and CC terms in the 249 DEGs specific to Pnoc receiver cells (boxed above). **L**) Examples of genes specifically altered in the S-R comparison of Pnoc receiver cells include *Oprl1*, *Arc*, and *Egr1*. (*p_adj_ < 0.05, Wilcoxon Rank Sum test, Benjamini-Hochberg correction)

Starting with the Crh pathway, we identified the putative receiving cells of Crh+ inhibitory neurons in the ACC, which were Crhr1-expressing L5 pyramidal neurons and, to a lesser extent, L2/3 neurons (**Figure 5E**), consistent with previous reports (Chen et al., 2020; Riad et al., 2022). To explore the transcriptional consequences of the predicted changes in Crh signaling, we took the Crhr1+ pyramidal neurons that were proximal (≤250µm) to Crh+ inhibitory neurons (**Figure 5F**) and performed differential expression between susceptible and resilient animals. Considering this analysis performed on any subset of L2/3 and L5 neurons may simply reproduce DEGs identified in the broader population of cells, we focused on DEGs unique to these proximal receiver cells to isolate DEGs putatively downstream of the predicted changes in Crh signaling. This analysis identified 127 DEGs, of which 69 (54.3%) were unique to Crh receiver cells and were not identified as DEGs in the broad population of L2/3 or L5 cells (**Figure 5G**). These DEGs were enriched for terms related to dendrites and synapses (**Figure 5G**), and include the Crhr1 gene itself, Crtc1, a transcriptionally-regulated gene whose dysfunction is associated with depression-like behavior (Cherix et al., 2022), and Snph, encoding a protein involved in activity-dependent mitochondrial transport in neurons (**Figure 5H**) (Park et al., 2016), all of which are upregulated in susceptible compared to resilient animals. These results indicate that reduced inhibitory Crh signaling in susceptible animals may be upstream of a compensatory upregulation of Crh receptors and, in parallel, upregulation of activity-dependent gene expression. We next determined which ligand-receptor pairs were driving the observed changes in predicted opioid signaling and found that predicted interactions were primarily observed between inhibitory neurons expressing genes encoding one of three opioid preproproteins (Pnoc, Penk, or Pdyn) and pyramidal neurons expressing Oprl1, encoding an opioid receptor (**Figure 5I**). We focused on Pnoc+ interneurons because Pnoc is processed into nociceptin, the primary ligand for the Oprl1 receptor (Henderson & McKnight, 1997). We isolated the Oprl1+ pyramidal cells proximal (<250µm) to Pnoc+ interneurons (**Figure 5J**) and performed differential expression between susceptible and resilient animals, again focusing on DEGs unique to this subpopulation of receiver cells. This analysis revealed 376 DEGs, of which 249 (66.2%) were unique to this group of receiver cells and were not found to be differentially expressed in any of the broader groups of pyramidal neurons (**Figure 5K**). The DEGs specific to the receiver cells were enriched for terms related to glutamatergic synapses and signaling (**Figure 5K**), and included the Oprl1 gene itself, as well as Arc and Egr1, which both encode immediate early genes that are transcribed in response to neuronal activation (**Figure 5L**) (Minatohara et al., 2016), all of which are upregulated in susceptible compared to resilient animals. Similar to what was observed for Crh, these findings imply that putatively reduced inhibitory opioid signaling in susceptible animals is upstream of a compensatory upregulation of opioid receptors alongside upregulation of activity-dependent immediate early gene transcripts.

### Enhanced neuron-astrocyte contact interactions are associated with increased susceptibility to stress-induced motivational impairment

Finally, we explored the finding that signaling strength between astrocytes and L2/3 pyramidal neurons was stronger in susceptible compared to resilient animals (**Figure 5B**). When we examined the specific signaling pathways altered between these cell types, we found Slitrk signaling, which plays a role in synapse regulation (Proenca et al., 2011), was elevated in the susceptible group (**Figure 6A**). Specifically, predicted cell adhesion interaction between astrocytic Slitrk2 and Ptprs or Ptprd (Ptprs/d) in L2/3 cells was increased in susceptible animals (**Figure 6B**). Taking the subset of Ptprs/d+ L2/3 cells proximal to Slitrk2+ astrocytes (**Figure 6C**), we performed differential expression between the susceptible and resilient groups and found that the majority of DEGs (52/58, 89.6%) were specific to these L2/3 receiver cells (**Figure 6D**). This set of DEGs was enriched for terms related to synapse organization (**Figure 6D**), and included Sncb, encoding synuclein beta which plays a role in vesicle endocytosis (Vargas et al., 2014), Stxbp1, encoding a syntaxin-binding protein with a role in vesicle exocytosis (Gerber et al., 2008), and Tbl1xr1, encoding a transcriptional regulator in which mutations have been associated with neurodevelopmental deficits in humans (Quan et al., 2020), all of which are upregulated in the resilient compared to susceptible group. These results suggest that increased Slitrk contact interactions between astrocytes and L2/3 pyramidal neurons are a feature of stress susceptibility which is associated with differences in synapse-related gene expression between susceptible and resilient mice.

**Figure 6.**
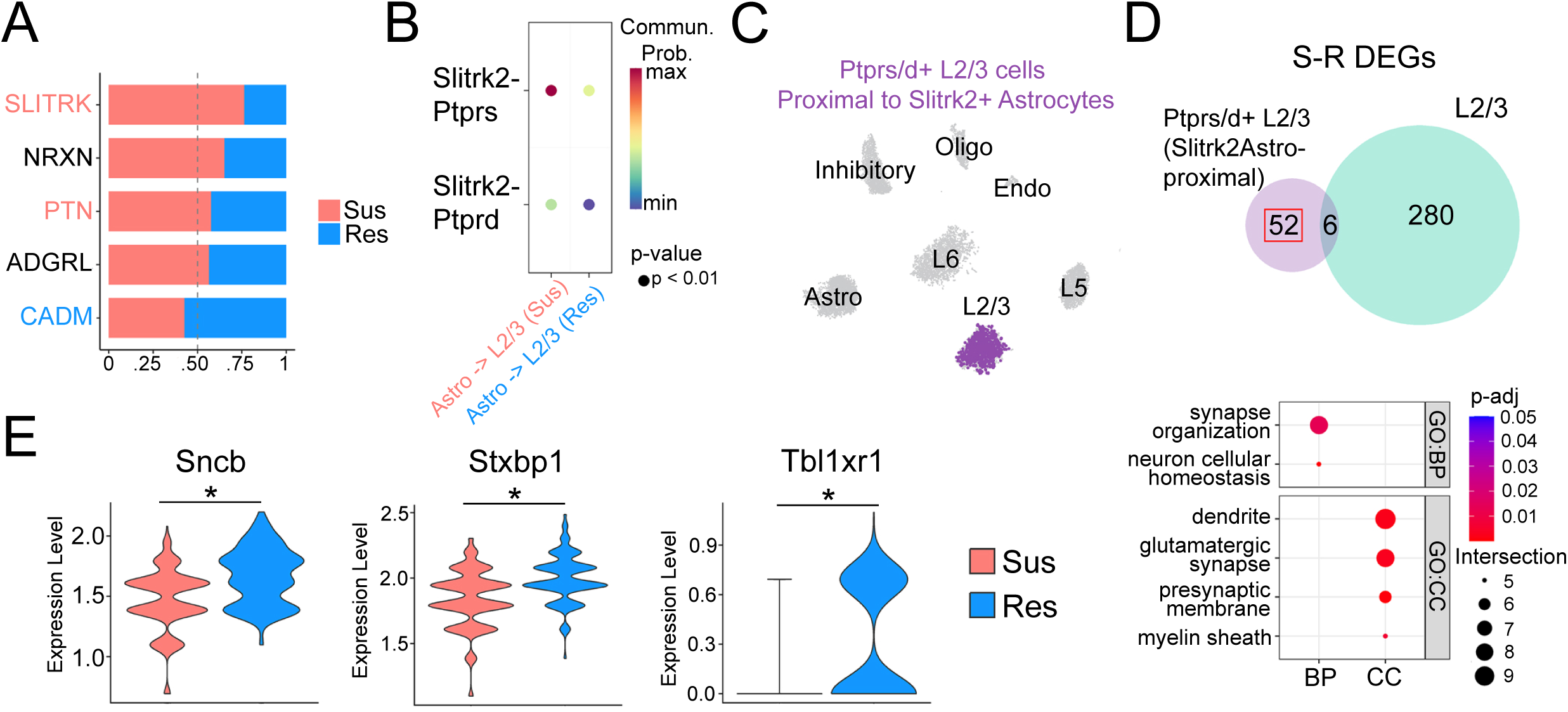
Stress susceptibility is marked by increased Slitrk signaling from astrocytes to L2/3 excitatory neurons. **A**) A stacked bar plot depicts the difference in relative signaling strength (information flow) between the susceptible and resilient conditions for specific signaling pathways with astrocytes as the source and L2/3 excitatory neurons as the targets (red and blue x-axis labels denote a statistically significant increase in signaling strength in the susceptible compared to the resilient condition and vice versa, respectively). **B**) A dotplot demonstrates the predicted Slitrk signaling from astrocytes to L2/3 excitatory neurons includes the Slitrk2-Ptprs and Slitrk2-Ptprd ligand-receptor pairs. **C**) A UMAP plot highlighting L2/3 cells that both 1) express *Ptprs* or *Ptprd* and 2) are within close proximity (55μm) of a *Slitrk2*+ astrocyte. **D**) A Venn diagram (top) depicting the overlap of susceptible vs. resilient (S-R) DEGs in the cells highlighted in **C** with the general populations of L2/3 neurons, along with a dot plot (bottom) showing the enrichment of gene ontology Biological Process (BP) and Cellular Component (CC) terms in the 52 DEGs specific to Slitrk receiver cells (boxed above). **E**) Examples of genes specifically altered in the S-R comparison of Slitrk receiver cells include *Sncb*, *Stxbp1*, and *Tbl1xr1*. (*p_adj_ < 0.05, Wilcoxon Rank Sum test, Benjamini-Hochberg correction)

## DISCUSSION

Here we describe adaptations in the ACC transcriptional landscape in animals with effortful reward seeking deficits following chronic stress exposure. The regulation of effort in pursuing rewards or goals is a key aspect of motivation and one that is impaired in neuropsychiatric conditions including depression. Classical measures of anhedonia in preclinical studies have largely focused on consummatory behaviors and preferences which may not reflect an accurate symptom profile in depression (Dichter et al., 2010). Amotivation is therefore a critical component and translationally valid domain to explore with improved assays (Spring et al., 2021). Our head-restrained effortful reinforcement task allows for this type of behavioral assessment and is designed to be compatible with in vivo physiology and optogenetic manipulation to facilitate future neural circuit characterizations. We identified unique patterns of differential gene expression in stress susceptibility and resilience using our novel paradigm as a readout of motivated behavior, combined with a spatial transcriptomic platform. This approach allowed us to preserve the anatomical and microcircuit environment of the ACC, a region critical for valuation, decision-making and goal-oriented behaviors, and gain greater insights into stress-induced genetic and molecular alterations with cell-level granularity.

Understanding the neurobiological bases for stress resilience has been of particular interest within the field with several genes, signaling pathways and physiological mechanisms identified as contributing factors across brain areas (Minerva et al., 2026; Bagot et al., 2016; Lorsch et al., 2019; Hing et al., 2024). These effects of stress have been characterized in whole brain regions using bulk and single-cell sequencing approaches. Our dataset reproduces several findings from earlier studies of cell type-specific transcriptional effects of stress on frontal cortical regions, including the sensitivity of excitatory neurons (Hing et al., 2024; Kwon et al., 2022) and astrocytes (Holt et al., 2024), as well as the central finding that stress alters the expression of synapse-related genes in excitatory neuron populations (Mallimo & Kusnecov, 2013; Jiang et al., 2024).

Leveraging the spatial dimension of our data, we extended these findings by identifying putative cell-cell signaling mechanisms upstream of these transcriptional changes. Overall, we find stress resilience is marked by increased communication strength between inhibitory and excitatory neurons, implicating excitatory-inhibitory balance in the ACC as an important feature of stress resilience. In particular, we homed in on altered Crh and opioid signaling between inhibitory and excitatory neurons, which were predicted to have increased signaling in stress resilient animals. Beyond its roles in the hypothalamus, Crh signaling has been implicated in behaviors related to cognition and reward (Bryce & Floresco, 2016; Hupalo, Bryce, et al., 2019; Hupalo, Martin, et al., 2019; Chen et al., 2020; Birnie et al., 2023). In addition, manipulation of prefrontal Crh interneurons has been shown to modulate both anxiety-related and motivated escape behaviors, with activation being associated with increased struggling and decreased avoidance and inhibition associated with the decreased struggling and increased avoidance (Chen et al., 2020). Similar to Crh, nociceptin opioid signaling is sensitive to stress (Devine et al., 2003; Witkin et al., 2014), and while little is known about local nociceptin signaling in the ACC, systemic administration of other Oprl1 agonists have been shown to have antidepressant- and anxiolytic-like effects in rodents (Gavioli & Calo’, 2006). Overall, the behavioral effects of altering Crh and nociceptin signaling are consistent with their roles in promoting a state of stress resilience. Notably, we observed increased expression of Crh and nociceptin receptors in susceptible animals, a common homeostatic signature of reduced signaling. Consistent with this, agonist-dependent receptor downregulation has been observed for Crhr1 (Kasagi et al., 2002) and Oprl1 (Hashimoto et al., 2002), and is thought to play a broad role in balancing synaptic signaling in the brain (Turrigiano, 2012). This putative reduction in inhibitory peptide signaling was associated with increased synapse-related and immediate early gene expression, indicating disinhibition of these excitatory populations in stress susceptible animals (**Figure 7**).

**Figure 7.**
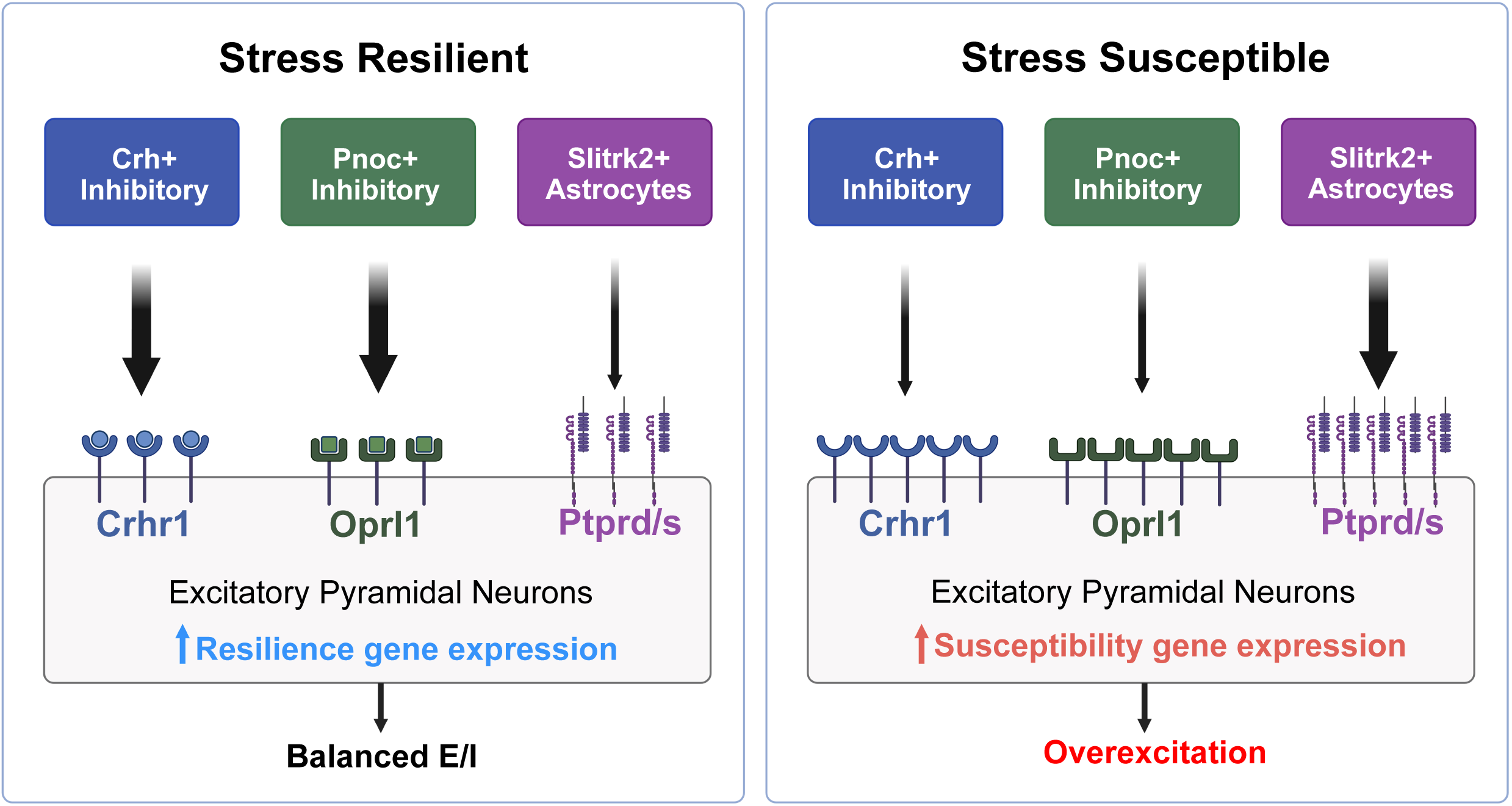
Working model of the cell-cell signaling mechanisms underlying transcriptomic and behavioral differences between stress resilient and stress susceptible animals in the ACC. In the resilient group (left), we observe a predicted increase in Crh and Pnoc inhibitory signaling and reduced astrocytic Slitrk2 signaling sent to excitatory cells which, we hypothesize, results in balanced excitation/inhibition (E/I) and a resilient transcriptional state. In the susceptible group (right), a reduction in Crh and Pnoc signaling (marked by high receptor expression) along with increased Slitrk2 signaling sent to excitatory cells underlies the susceptible transcriptional state and, we hypothesize, leads to overexcitation in the ACC.

In addition to neuronal signaling, we observed stress-mediated changes in gene expression and cell-cell communication in astrocytes. Astrocytes are known to be both sensitive to chronic stress (Mayhew et al., 2015) and to play an important role in mediating stress resilience (Holt et al., 2024) and in driving antidepressant behavioral effects in mice (Yue et al., 2025). Moreover, bidirectional crosstalk between neurons and astrocytes in the lateral habenula mediates stress-induced depressive-like behaviors (Xin et al., 2025). Consistent with this, we find that stress elevates levels of Ptgds in both resilient and susceptible animals, indicating the induction of astrocyte reactivity (Matusova et al., 2023). However, in resilient animals, we observe simultaneous upregulation of the gene encoding calcineurin, which has been shown to have anti-inflammatory effects in reactive astrocytes (Fernandez et al., 2007) potentially blunting the effects of astrocyte reactivity in resilient animals. Using spatial information to identify astrocytic cell-cell signaling interactions that may be altered by stress, we found that astrocytes show elevated communication strength with L2/3 pyramidal neurons in the susceptible condition, which is primarily driven by elevated Slitrk2 signaling. Interestingly, Slitrk proteins have been implicated in isoform-specific effects on balancing inhibitory and excitatory synapses (Yim et al., 2013), with Slitrk2-Ptprs interactions being involved in excitatory synapse formation. Increased Slitrk2 signaling in stress susceptible animals may therefore further contribute to increased excitability of ACC pyramidal neurons in the susceptible condition (**Figure 7**).

Synthesizing the results from the cell-cell communication analyses, our findings indicate that stress susceptibility is associated with increased ACC excitability (**Figure 7**). Clinical studies link overactivity of the subgenual cingulate cortex with major depressive disorder (Hamani et al., 2011). Modulating activity in this region via antidepressant drugs such as ketamine (Alexander et al., 2021) or by deep brain stimulation (Mayberg et al., 2005; Lozano et al., 2008; Fujimoto et al., 2024), is associated with treatment efficacy. In preclinical studies, the association between stress and medial prefrontal excitability has previously been implicated in anhedonia in mice (Ferenczi et al., 2016; Stanton et al., 2019) and nonhuman primates (Alexander et al., 2019). It should be noted that diminished ACC function has also been associated with motivational deficits related to reduced voluntary speech and actions (Devinsky et al., 1995). Importantly, the ACC is a large structure and anatomical and functional segregation of subregions may differ between humans and rodents (van Heukelum et al., 2020). Additionally, brain networks including the ACC may be similarly impaired by hyper- or hypoactivity. In line with this, we found that stress resilience was not simply marked by the absence of susceptibilty-associated transcriptional and signaling changes, but instead by a distinct set of changes associated with inhibition that may serve to temper stress-induced increases in excitability. Future studies should aim to manipulate Crh, nociceptin, and Slitrk2 signaling in the ACC of mice subject to chronic social defeat stress to clarify their precise role in mediating stress-induced changes in motivated reward behaviors. Future studies should also examine the circuit mechanisms mediating stress resilience and susceptibility to connect the observed transcriptional and signaling changes with altered activity of specific projections to and from the ACC.

## Acknowledgements

P.K.P conceptualized the study with input from M.K and C.L. P.K.P and M.K collected data and samples. P.K.P and D.R analyzed data and prepared the manuscript with input from C.L. J.R contributed to the design and construction of behavioral equipment H.M. assisted with pilot sequencing experiments. We would like to thank the Weill Cornell Medicine Epigenomics Core Facility for guidance on experiment planning, and for conducting sequencing and cDNA library preparation. We would also like to thank Jolin Chou and Shawn Singh for technical assistance. This work was supported by the following funding sources: NIH MH127291 and WCM JumpStart Research Career Development awards to P.K.P; NIH MH142082 award to D.R; NIH DA047851, MH109685, MH118451, Hope for Depression Research Foundation, and Rita Allen Foundation awards to C.L.

