## Supplementary Information for "Stress-induced motivational impairment is marked by diminished frontocortical cellular communication and neuropeptide signaling"

**Description of Supporting Information Figures & Tables:**

**Supplementary Figures**

**Supplementary Figure 1.** The CNSDS protocol results in attacks directed to male and female mice and alterations in anxiety-related behavior.

**Supplementary Figure 2.** No correlation between stress-induced changes in reward seeking and social interaction or anxiety-related behavior in resilient and susceptible groups.

**Supplementary Figure 3.** Analysis of spot-level spatial transcriptomics data reveal laminar spot clusters and limited expression differences between groups.

**Supplementary Figure 4.** Spatial deconvolution yields data with single-cell resolution.

**Supplementary Figure 5.** Differentially expressed genes consistent gene expression profiles across individual replicates.

**Supplementary Figure 6.** The degree of predicted inhibitory to excitatory signaling in controls lies between the susceptible and resilient groups.

**Supplementary Tables**

**Supplementary Table 1.** Table of differentially expressed genes identified in the spot-level analysis (supplied as a separate excel file)

**Supplementary Table 2.** Table of differentially expressed genes identified in the cell-level analysis (supplied as a separate excel file)

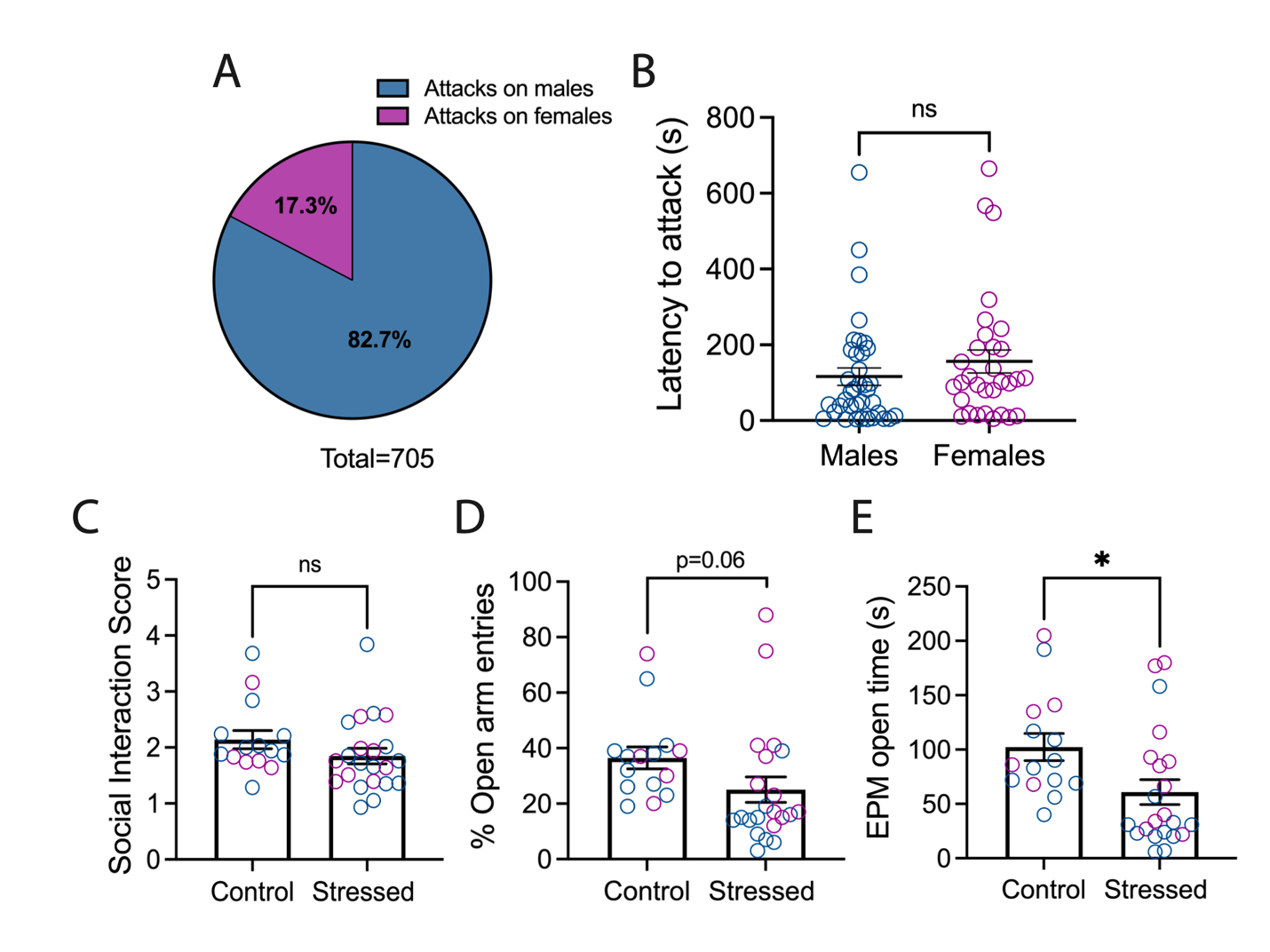

**Supplementary Figure 1. The CNSDS protocol results in attacks directed to male and female mice and alterations in anxiety-related behavior. A)** Proportion of attacks on males and females across 10 days and **B**) latency to attack the male and female test mouse during each defeat session (n=4 males, 2 females). **C**) Social interaction scores for control and stressed animals in the open field social interaction (OFSI) assay (Unpaired t test; p=0.1779). **D**) Percent open arm entries in the elevated plus maze (EPM) for control and stressed animals (Unpaired t test; p=0.0664). **E**) Time spent in the open arms of the EPM for control and stressed mice (Unpaired t test; p=0.0198; n=15 control, n=22 stressed; *p<0.05).

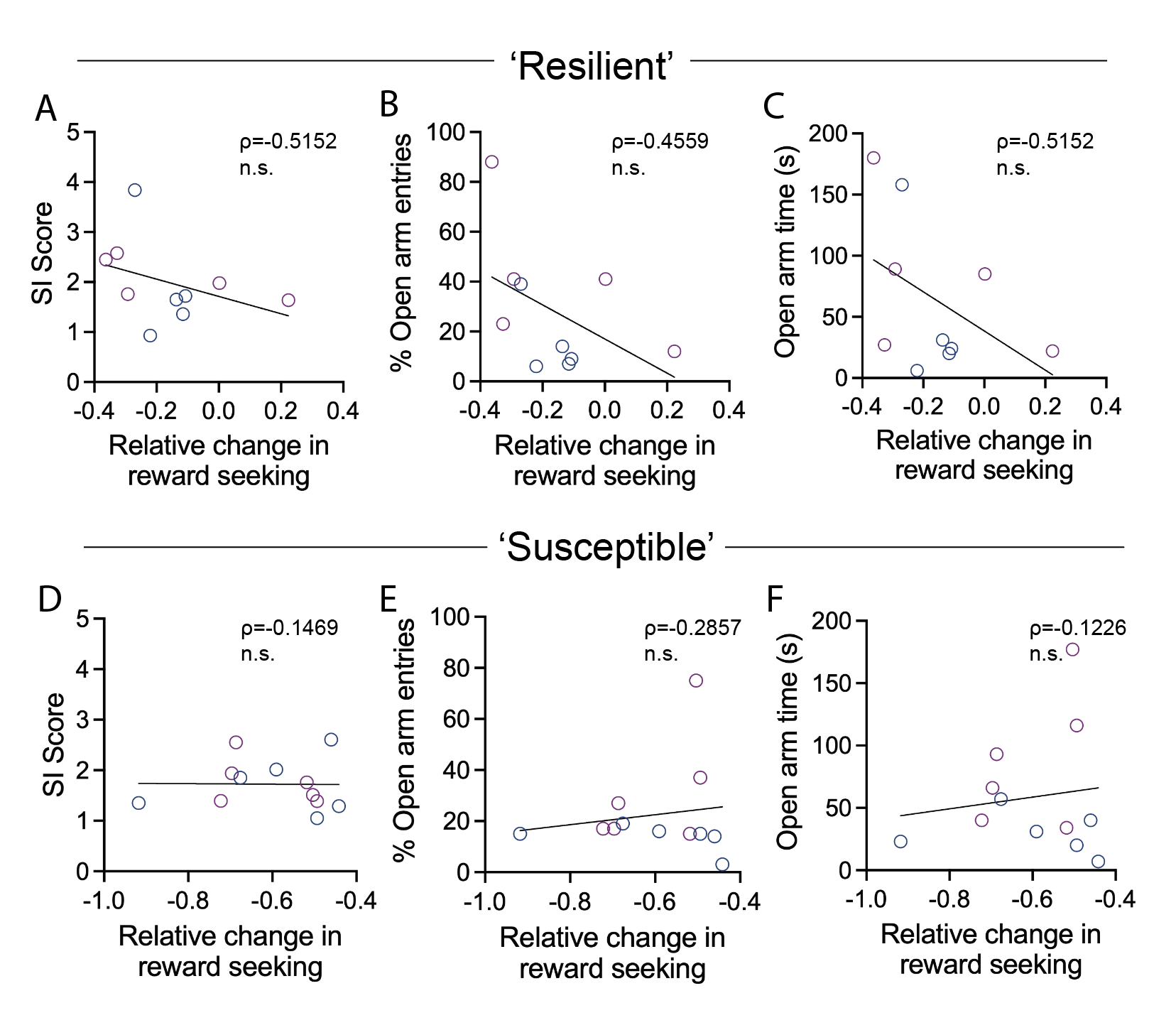

**Supplementary Figure 2. No correlation between stress-induced changes in reward seeking and social interaction or anxiety-related behavior in resilient and susceptible groups.**  **A-C)** Scatterplots indicating no significant correlation between relative change in reward seeking in the Pavlovian task and social interaction score (SI) or open arm entries and time in the elevated plus maze in mice classified as resilient to reward deficits (n=10 mice; blue symbols = males; magenta symbols = females). **D-F**) Scatterplots indicating no significant correlation between relative change in reward seeking in the Pavlovian task and social interaction score (SI) or open arm entries and time in the elevated plus maze in mice classified as susceptible to reward deficits (n=12 mice). Spearman’s correlation.

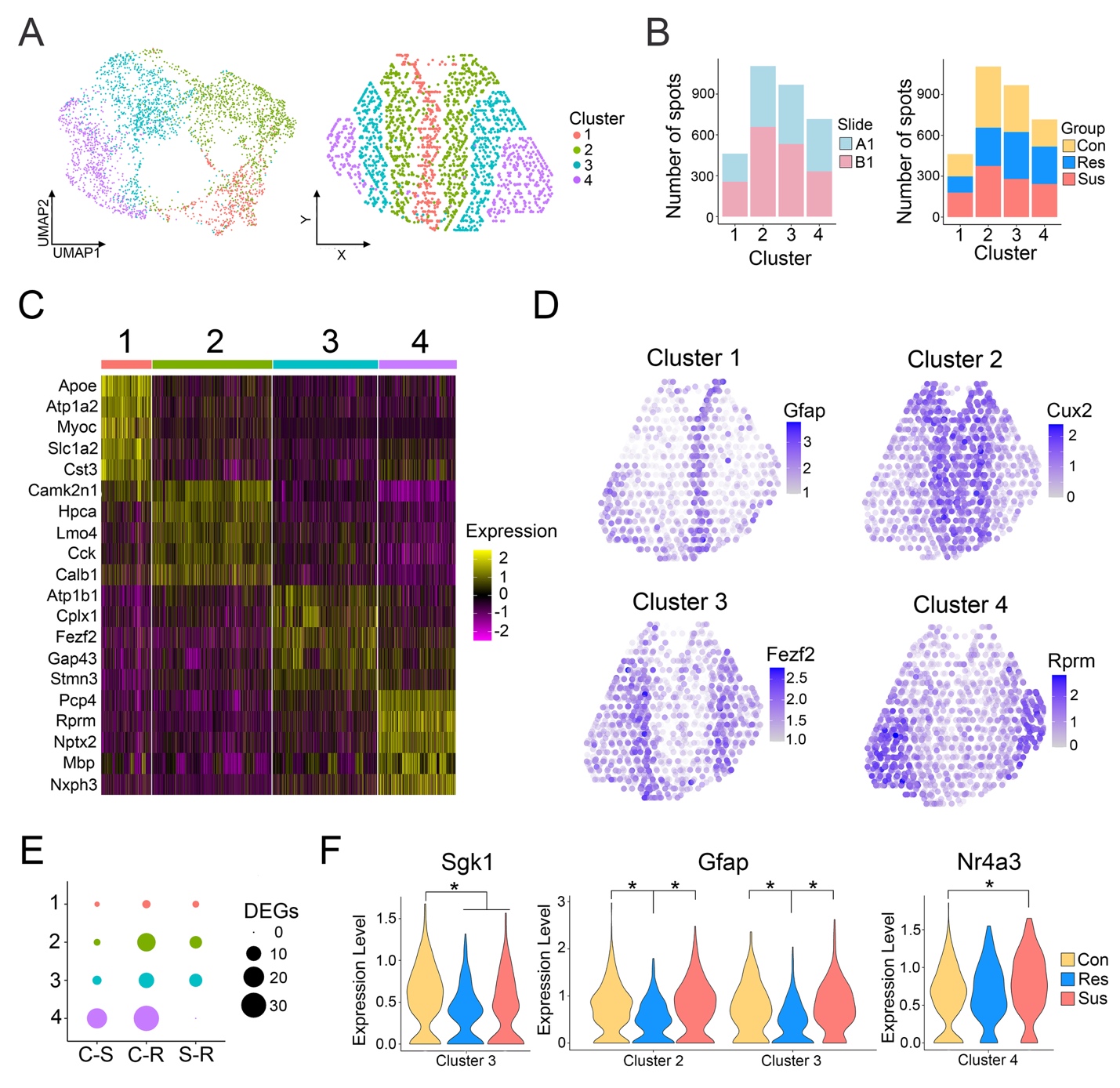

**Supplementary Figure 3.** **Analysis of spot-level spatial transcriptomics data reveal laminar spot clusters and limited expression differences between groups. A**) UMAP plot (left) and consensus spatial plot (right) of spot-level data. **B**) Stacked bar plots demonstrate a similar proportion of spot clusters were identified across Visium slides (left) and groups (right). **C**) A heatmap depicting the top 5 marker genes for each cluster. **D**) Spatial feature plots for one of the marker genes shown in **C** for each cluster. **E**) A dotplot shows differentially expressed genes (DEGs) within each spot cluster for the control vs. susceptible (C-S), control vs. resilient (C-R), and susceptible vs. resilient (S-R) comparisons. **F**) Example DEGs include *Sgk1* in cluster 3, *Gfap* in clusters 2 & 3, and *Nr4a3* in cluster 4.(*: p_adj_ < 0.05, Wilcoxon Rank Sum test, Benjamini-Hochberg correction)

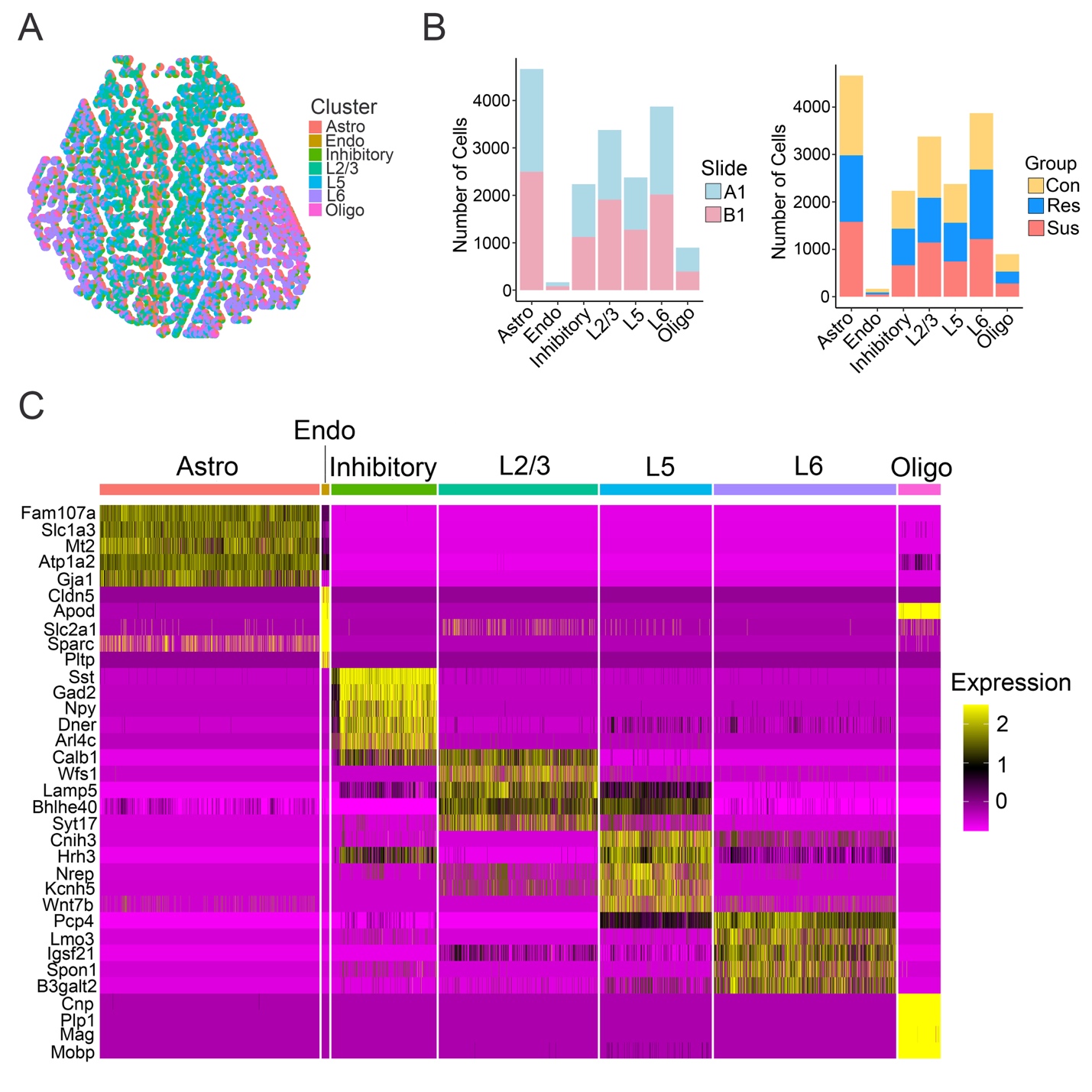

**Supplementary Figure 4. Spatial deconvolution yields data with single-cell resolution. A**) A spatial consensus pie plot in which each Visium spot is represented by a pie chart that depicts the proportions of cell types within that spot identified during deconvolution. **B**) Stacked bar plots demonstrating that, after deconvolution, a similar proportion of cell types were identified across Visium slides (left) and groups (right). **C**) A heatmap depicting the top 5 marker genes for each cell cluster.

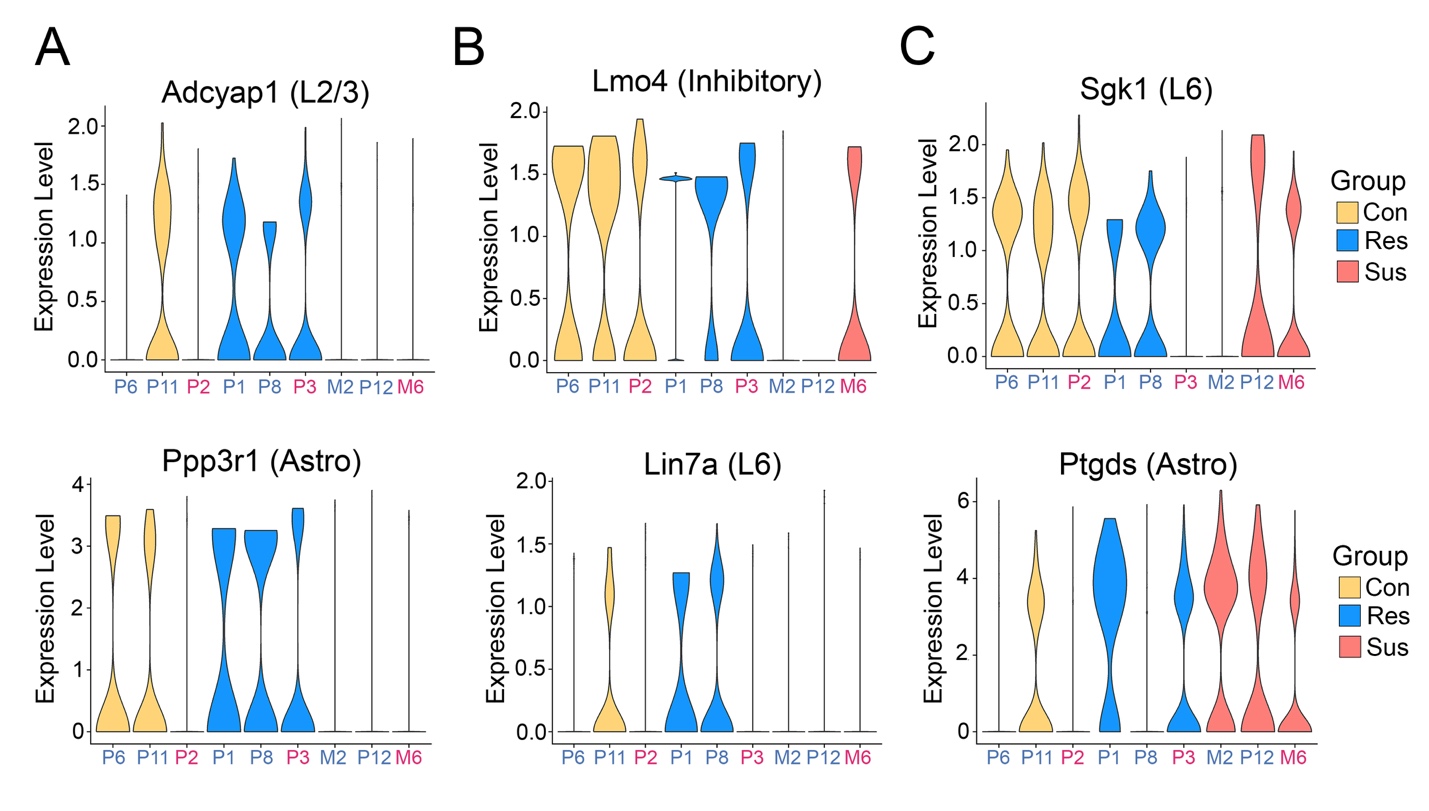

**Supplementary Figure 5. Differentially expressed genes consistent gene expression profiles across individual replicates.** Violin plots separated by replicate are shown for the resilience genes depicted in **Figure 4C** (**A**), the susceptibility genes depicted in **Figure 4D** (**B**), and the stress genes depicted in **Figure 4E** (**C**). X-axis text is colored by the animal’s sex (blue: male; red: female)

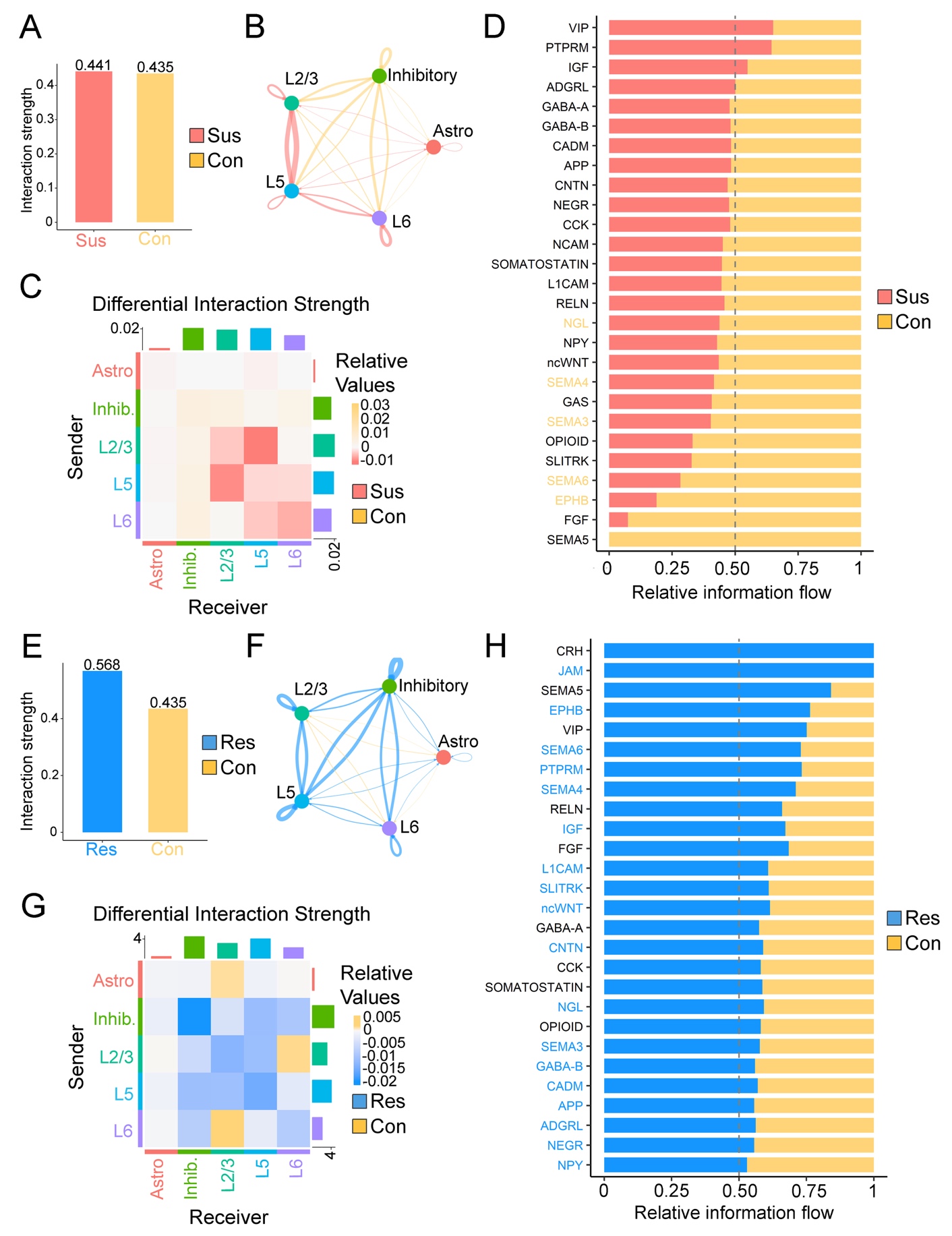
**Supplementary Figure 6. The degree of predicted inhibitory to excitatory signaling in controls lies between the susceptible and resilient groups.** **A**) A barplot shows the difference in overall interaction strength between the susceptible and control groups. The differential interaction strength between specific cell clusters is shown as a circle plot (**B**) and as a heatmap (**C**), with red and yellow representing interactions with increased probability in the susceptible condition (compared to control) or vice versa, respectively. Edge width and color intensity represent the magnitude of difference between groups in the circle plot and heatmap, respectively. **D**) A stacked bar plot depicts the difference in relative signaling strength (information flow) between the susceptible and control conditions for specific signaling pathways with inhibitory neurons as the source and excitatory neurons as the targets (yellow x-axis labels denote a statistically significant increase in signaling strength in the control compared to susceptible condition). **E**) A barplot shows the difference in overall interaction strength between the resilient and control groups. The differential interaction strength between specific cell clusters is shown as a circle plot (**F**) and as a heatmap (**G**), with blue and yellow representing interactions with increased probability in the resilient condition (compared to control) or vice versa, respectively. Edge width and color intensity represent the magnitude of difference between groups in the circle plot and heatmap, respectively. **H**) A stacked bar plot depicts the difference in relative signaling strength (information flow) between the susceptible and control conditions for specific signaling pathways with inhibitory neurons as the source and excitatory neurons as the targets (blue x-axis labels denote a statistically significant increase in signaling strength in the resilient compared to control condition).
